# La-related protein 4 is enriched in vaccinia virus factories and is required for efficient viral replication in primary human fibroblasts

**DOI:** 10.1101/2023.03.10.532125

**Authors:** Pragyesh Dhungel, Djamal Brahim Belhaouari, Zhilong Yang

## Abstract

In addition to the 3’-poly(A) tail, vaccinia virus mRNAs synthesized after viral DNA replication (post-replicative mRNAs) possess a 5’-poly(A) leader that confers a translational advantage in virally infected cells. These mRNAs are synthesized in viral factories, the cytoplasmic compartment where vaccinia virus DNA replication, mRNA synthesis, and translation occur. However, a previous study indicates that the poly(A)-binding protein (PABPC1)-which has a well-established role in RNA stability and translation-is not present in the viral factories. This prompts the question of whether another poly(A)-binding protein engages vaccinia virus post-replicative mRNA in viral factories. In this study, we found that La-related protein 4 (LARP4), a poly(A) binding protein, was enriched in viral factories in multiple types of cells during vaccinia virus infection. Further studies showed that LARP4 enrichment in the viral factories required viral post-replicative gene expression and functional decapping enzymes encoded by vaccinia virus. We further showed that knockdown of LARP4 expression in human foreskin fibroblasts (HFFs) significantly reduced vaccinia virus post-replicative gene expression and viral replication. Interestingly, the knockdown of LARP4 expression also reduced 5’-poly(A) leader-mediated mRNA translation in vaccinia virus-infected and uninfected HFFs. Together, our results identified a poly(A)-binding protein, LARP4, enriched in the vaccinia virus viral factories and facilitates viral replication and mRNA translation.

**Importance:** Poxviruses are a family of large DNA viruses comprising members infecting a broad range of hosts, including many animals and humans. Poxvirus infections can cause deadly diseases in humans and animals. Vaccinia virus, the prototype poxvirus, encodes over 200 open reading frames (ORFs). Over 90 of vaccinia virus ORFs are transcribed post-viral DNA replication. All these mRNAs contain a 5’-poly(A) leader, as well as a 3’-poly(A) tail. They are synthesized in viral factories, where vaccinia virus DNA replication, mRNA synthesis and translation occur. However, surprisingly, the poly(A) binding protein (PABPC1) that is important for mRNA metabolism and translation is not present in the viral factories, suggesting other poly(A) binding protein(s) may be present in viral factories. Here we found another poly(A)-binding protein, La-related protein 4 (LARP4), is enriched in viral factories during vaccinia virus infection. We also showed that LARP4 enrichment in the viral factories depends on viral post-replicative gene expression and functional viral decapping enzymes. The knockdown of LARP4 expression in human foreskin fibroblasts (HFFs) significantly reduced vaccinia virus post-replicative gene expression and viral replication. Overall, this study identified a poly(A)-binding protein that plays an important role in vaccinia virus replication.

## Introduction

Poxviruses are a large family of double-stranded DNA viruses causing many significant diseases in humans and animals, including smallpox, one of the most notorious infectious diseases in human history [1]. Mpox virus, the causative agent of mpox (previously named as monkeypox), is endemic in western and central Africa and, with an increasing presence in other regions of the world, poses a growing global threat to public health [2–4]. Vaccinia virus (VACV) is the prototype poxvirus and is a highly relevant surrogate to study high pathogenic poxviruses (e.g., mpox and smallpox viruses) due to their high similarity with >95% genome identity [5]. VACV is also the vaccine against smallpox and mpox. VACV encodes over 200 open reading frames (ORFs) in its ~200kbp genomes that are expressed in a cascade manner [6]. While the early genes (118 ORFs) are expressed before viral DNA replication, the intermediate and late genes (~90 ORFs) are expressed after viral DNA replication [7–9]. These intermediate and late genes are collectively termed post-replicative genes.

All the post-replicative mRNAs have a 5’-poly(A) leader, in addition to a 3’-poly(A) tail, with heterogenous lengths that are likely generated by polymerase transcription slippage on a triple-thymine stretch of the template strand of the DNA promoter [10, 11]. The role of 3’-poly(A) tail in mRNA metabolism and translation is well established with countless publications. Studies from us and others indicate that the 5’-poly(A) confers a translational advantage in poxvirus-infected cells undergoing host shutoff [12]. Our studies also indicate that the 5’-poly(A) is a cap-independent translation enhancer, which is promoted in VACV infection in a virally encoded decapping enzyme-dependent manner [13–x16]. RACK1, a 40S ribosomal protein, is specifically required for poly(A)-leader mediated translation and depends on RACK1 phosphorylation status during VACV infection [16]. ZNF598, a quality control sensor of collided ribosomes, is required for the translation of VACV mRNAs, including the poly(A)-headed intermediate and late mRNAs and early mRNAs that mostly do not harbor a 5’-poly(A) leader [17, 18]. However, interestingly, the well-known poly(A)-binding protein, PABPC1, is located outside of VACV viral factories [19], where viral mRNAs are produced and translated [20]. Given that poxvirus post-replicative mRNAs have a 5’-poly(A) leader and 3’-poly(A) tail, it is intriguing to know if any cellular poly(A)-binding proteins are recruited by poxviruses for its replication.

RNA-binding proteins play diverse roles in regulating RNA metabolism and translation. One such family of RNA-binding proteins is the La protein family that comprises the La RNA-binding motifs and the RNA recognition motif [21]. As a member of the La protein family, La-related protein 4 (LARP4) is known to bind to poly(A) RNA sequences (e.g., the 3’-poly(A) tail), the receptor for activated C kinase 1 (RACK1), and PABPC1 [22, 23]. LARP4 promotes mRNA stability by binding to the 3’ poly(A) tail and preventing RNA deadenylation. The RNA-binding La motif, RNA recognition motif, and poly(A) binding protein-interacting domain in LARP4 are necessary for its mRNA stability enhancement function. Moreover, the predominantly cytoplasmic LARP4 is also postulated to play a role in translation as it is shown to associates with actively translating ribosomes, although the precise function is still unclear [21–23].

In the present study, with the objective to examine the roles of poly(A)-binding proteins in VACV infection, we found that LARP4 was enriched in “virus factories,” a distinct cytoplasmic compartment where viral genome replication, intermediate and late transcription, and mRNA translation occur during VACV infection [20]. The recruitment of LARP4 to viral factories requires VACV post-replicative gene expression and functional viral decapping enzymes. We further demonstrated that the knockdown of LARP4 was unfavorable for VACV replication and inhibited VACV replication at the post-DNA replication stage in primary human foreskin fibroblasts (HFFs). Moreover, we found that the knockdown of LARP4 significantly decreased 5’-poly(A) leader-mediated mRNA translation. Our results identified the poly(A)-binding protein LARP4 as a pro-VACV replication host factor, which is required to translate mRNA with a 5’-poly(A) leader efficiently in primary human fibroblasts.

## Results

### LARP4 is enriched in viral factories during VACV infection

VACV replicates in the cytoplasm forming cytoplasmic DNA regions known as viral factories, where transcription and translation of viral intermediate and late mRNA occur [20]. Using confocal immunofluorescence microscopy, we observed that a well-known poly(A)-binding protein, PABPC1, that plays a significant role in mRNA translation and stability by binding to poly(A) stretch of mRNA-, was not enriched in, but in fact, excluded from viral factories in VACV-infected HFFs (**Fig. 1AC**). This result is consistent with a previous report that PABPC1 is distributed outside the viral factories in VACV-infected normal human dermal fibroblasts [19]. Interestingly, another poly(A) binding protein, LARP4, was enriched in viral factories during VACV infection of HFFs, while it was dispersed in the cytoplasm of uninfected HFFs (**Fig. 1BC).** The endogenous LARP4 enrichment in viral factories was detected as early as 4 hpi (**Fig. 1D**). We also observed the enrichment of LARP4 in viral factories in VACV-infected A549 cells and HeLa Cells (**Fig. 2ABC**). We did not observe a consistent pattern of increased LARP4 protein levels by Western blotting analysis during VACV infection (data not shown), suggesting the main effect of VACV infection is to re-localize and enrich LARP4 to viral factories.

**Figure 1.**
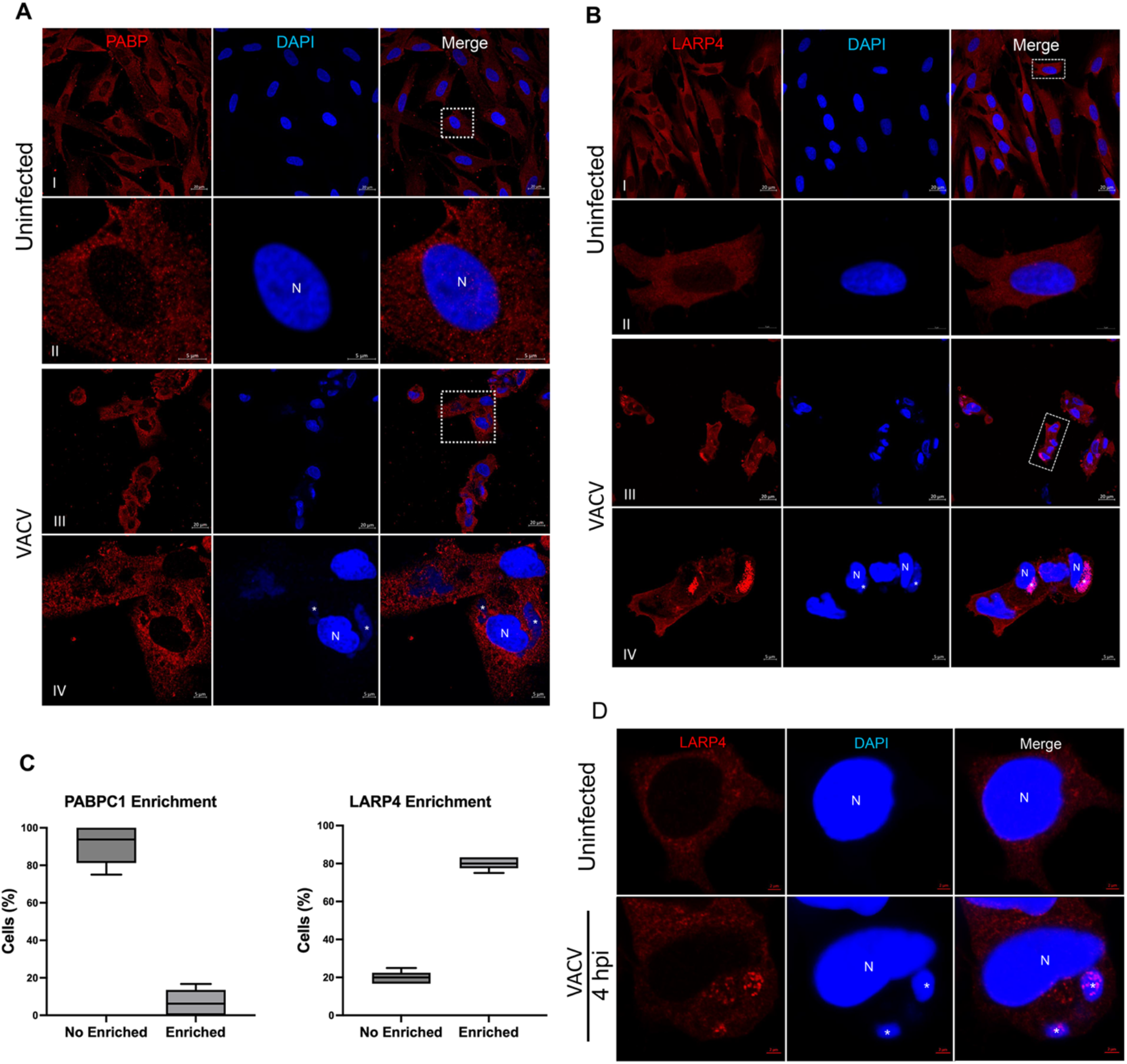
Enrichment of LARP4 in viral factories during VACV Infection of HFFs. (A) Confocal immunofluorescence microscopy of uninfected (I, II) and VACV-infected HFFs (III, IV) (MOI=2, 12 hpi). Anti-PABPC1 was used to determine the localization of PABPC1 (red), and DAPI (blue) was utilized to visualize cellular nuclei (N) and the viral factories (indicated by *). (High-magnification images of the cell boxed in (I) and the infected boxed cells in (III) are shown in (II) and (IV), respectively. The presented images are representative images of multiple fields of view. **(B)** Confocal immunofluorescence microscopy of uninfected (I, II) and VACV-infected HFFs (III, IV) (MOI=2, 12 hpi). Anti-LARP4 was used to determine the localization of LARP4 (red), while DAPI staining was used to visualize cellular nuclei (N) and the viral factories (indicated by *). High-magnification images of the cell boxed in (I) and the infected boxed cells in (III) are shown in (II) and (IV), respectively. The scale bars are 5 μm and 20 μm for high and low magnifications, respectively. **(C)** Quantification of the percentages of cells with PABPC1 and LARP4 enrichment in viral factories during VACV infection, a count of at least 25 cells was performed in different random confocal microscopy views. The resulting graph represents the percentage of cells with PABPC1 and LARP4 enrichment in viral factories. **(D)** Confocal immunofluorescence microscopy of uninfected and VACV-infected HFFs (MOI=2). At 4 hpi, HFFs were fixed and stained with indicated antibodies or DAPI. Anti-LARP4 was used to determine the localization of LARP4 (red), and DAPI (blue) was used to stain cellular nuclei (N) and viral factories (indicated by *). The scale bars are 5 μm.

**Figure 2.**
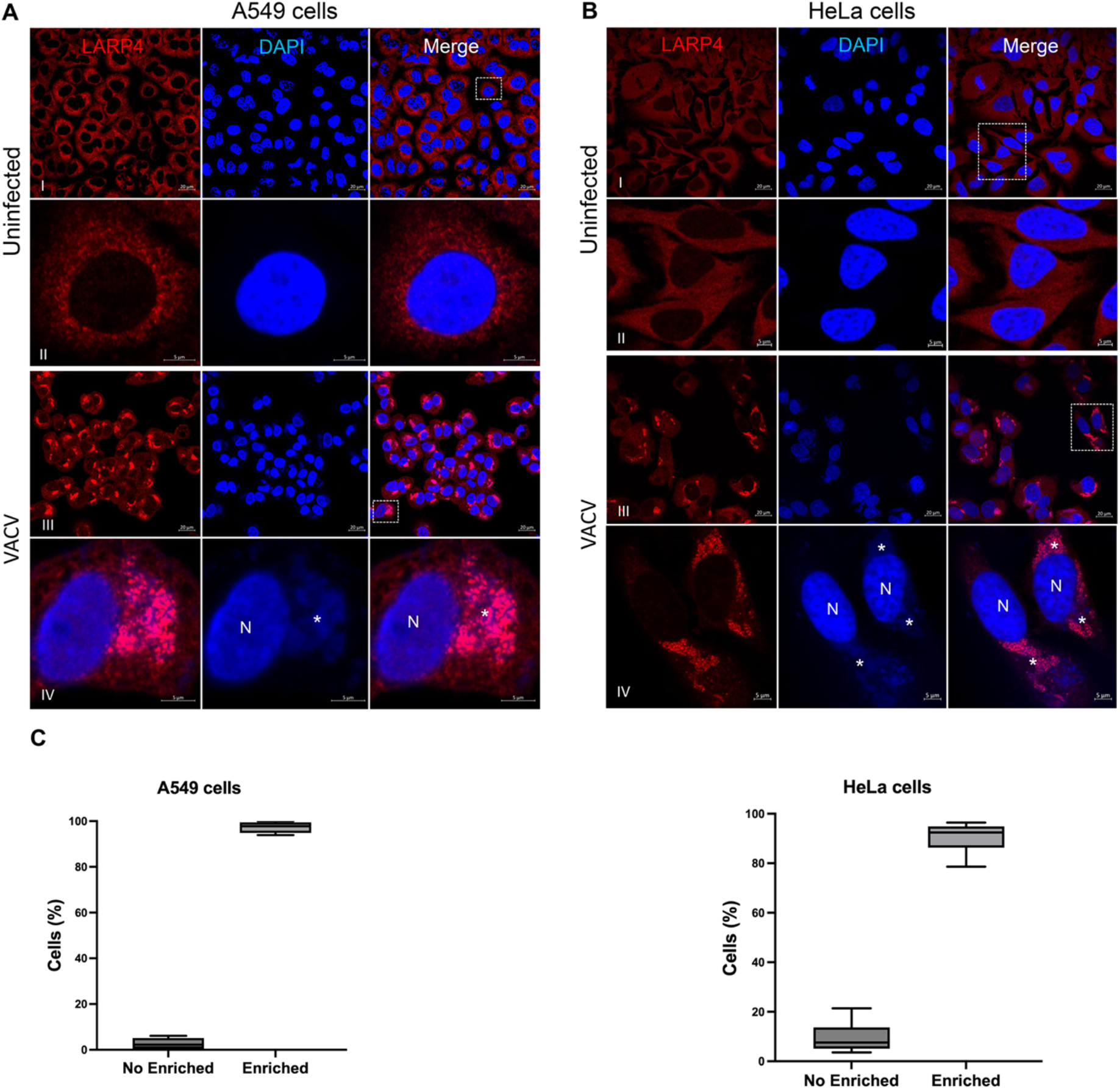
Enrichment of LARP4 in viral factories in VACV-infected A549 cells and HeLa cells. **(A)** Confocal immunofluorescence microscopy of uninfected (I, II) and VACV-infected A549 Cells (III, IV) (MOI=2, 12 hpi). Anti-LARP4 was used to determine the localization of LARP4 (red), and DAPI (blue) was used to stain cellular nuclei (N) and viral factories (indicated by *). (II) shows a high magnification of the cell boxed in (I), and (IV) shows a high magnification of the infected boxed cells in (III). **(B)** Confocal immunofluorescence microscopy of uninfected (I, II) and VACV-infected HeLa cells (III, IV) (MOI=2, 12 hpi). At 12 hpi, HeLa cells were fixed and stained with indicated antibodies. Anti-LARP4 was used to determine the localization of LARP4 (red), and DAPI (blue) was used to stain cellular nuclei (N) and viral factories (indicated by *). (II) shows a high magnification of the cell boxed in (I), and (IV) shows a high magnification of the infected boxed cells in (III). In A and B, the presented images are representatives of multiple fields of view. The scale bars are 5 μm and 20 μm for high and low magnifications, respectively. **(C)** The graphs show the quantification of cells with LARP4 enrichment in viral factories of VACV-infected A549 and HeLa cells. The percentage of cells with LARP4 enrichment in viral factories was counted across multiple random confocal microscopy views, with a total count of at least 160 cells per sample.

### Enrichment of LARP4 in viral factories requires viral post-DNA replication events and functional VACV decapping enzymes

VACV viral factories are formed after viral genomic DNA replication [20]. To further investigate if VACV viral DNA replication alone without intermediate gene transcription and following events is sufficient for LARP4 enrichment, we utilized a recombinant VACV (vΔA23) with the ORF encoding intermediate transcription factor gene A23 deleted [24]. While this virus undergoes genomic DNA replication and form viral factories, the post-replicative gene expression is blocked at the step of intermediate transcription, and no intermediate and late mRNAs are produced [24]. HFFs were infected with vΔA23, and subcellular localization of LARP4 was determined by confocal microscopy. At 12 hpi, we observed many viral factories. However, LARP4 was not enriched but excluded from the factories in HFFs and HeLa cells (**Fig. 3ABC**), indicating the requirement of post-DNA replicative event(s) to recruit LARP4 to viral factories.

**Figure 3.**
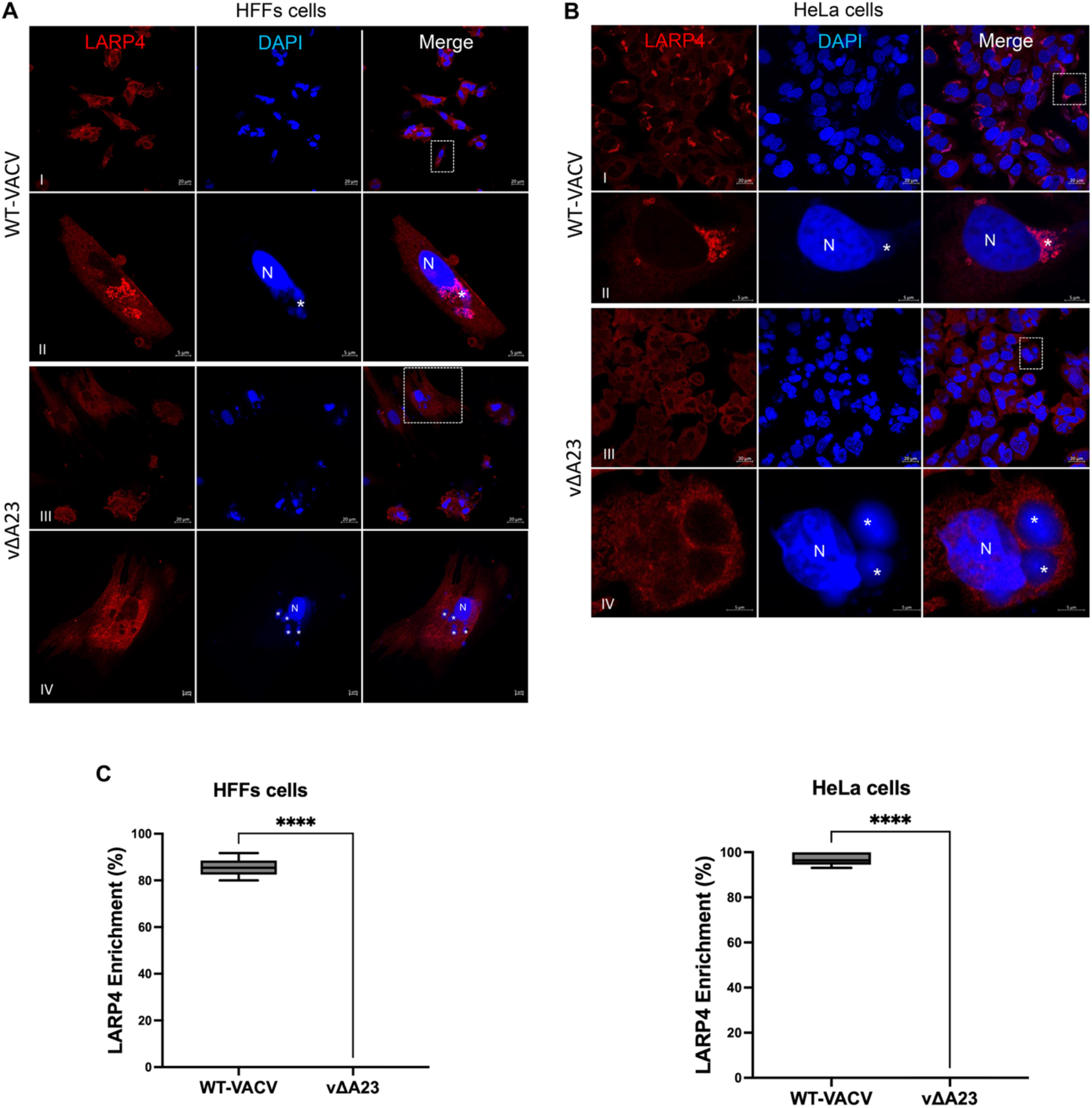
LARP4 Enrichment in viral factories requires post-replicative gene expression. **(A)** Confocal immunofluorescence microscopy of WT-VACV infected HFFs (I, II) and vΔA23 HFFs (III, IV) (MOI=2, 12 hpi). Anti-LARP4 was used to determine the localization of LARP4 (red), and DAPI (blue) was used to stain cellular nuclei (N) and viral factories (indicated by *). (II) shows a high magnification of the cell boxed in (I), and (IV) shows a high magnification of the cells boxed in (III). **(B)** Confocal immunofluorescence microscopy of WT-VACV infected HeLa cells (I, II) and vΔA23 infected HeLa cells (III, IV) (MOI=2, 12 hpi). Anti-LARP4 was used to determine the localization of LARP4 (red), and DAPI (blue) was used to stain cellular nuclei (N) and viral factories (indicated by *). (II) shows a high magnification of the cell boxed in (I), and (IV) shows a high magnification of the cells boxed in (III). The presented images are representatives of multiple fields of view. The scale bars are 5 μm and 20 μm for high and low magnifications, respectively. **(C)** Comparative analysis of LARP4 enrichment in viral factories of HFFs (left) and HeLa (right) infected with WT-VACV or vΔA23. P-values were obtained using the *Student’s t-test*, and the graph shows that LARP4 enrichment is significantly lower (P<0.01) in cells infected with vΔA23 than in cells infected with WT-VACV.

Next, we used another recombinant VACV, vD9muD10mu, with both decapping enzymes (D9 and D10) inactivated by mutating their Nudix motifs that harbor their decapping enzyme activity. The mutations in D9 and D10 render them losing the ability to decap cellular and viral mRNAs to speed up RNA decay [25]. VACV intermediate and late mRNAs are still efficiently produced in vD9muD10mu-infected cells [14, 15, 25, 26]. To examine if LARP4 is enriched in viral factories during vD9muD10mu infection, we used the A549DKO cell line that allows VACV replication by knocking out two antiviral genes encoding PKR and RNase L. The knockout of RNase L also ensures intact cellular and viral mRNAs in vD9muD10mu infection, as VACV could activate RNase L by viral dsRNA for RNA cleavage. In A549DKO cells, viral intermediate and late proteins are also produced, though at lower levels [26]. Interestingly, our results revealed that LARP4 was minimally enriched in the viral factories of vD9muD10mu-infected A549 cells or A549DKO comparing to WT-VACV infected cells under same imaging specifications (**Fig. 4ABC**). The results suggests that functional decapping activities are necessary for LARP4 recruitment to the viral factories during VACV infection.

**Figure 4.**
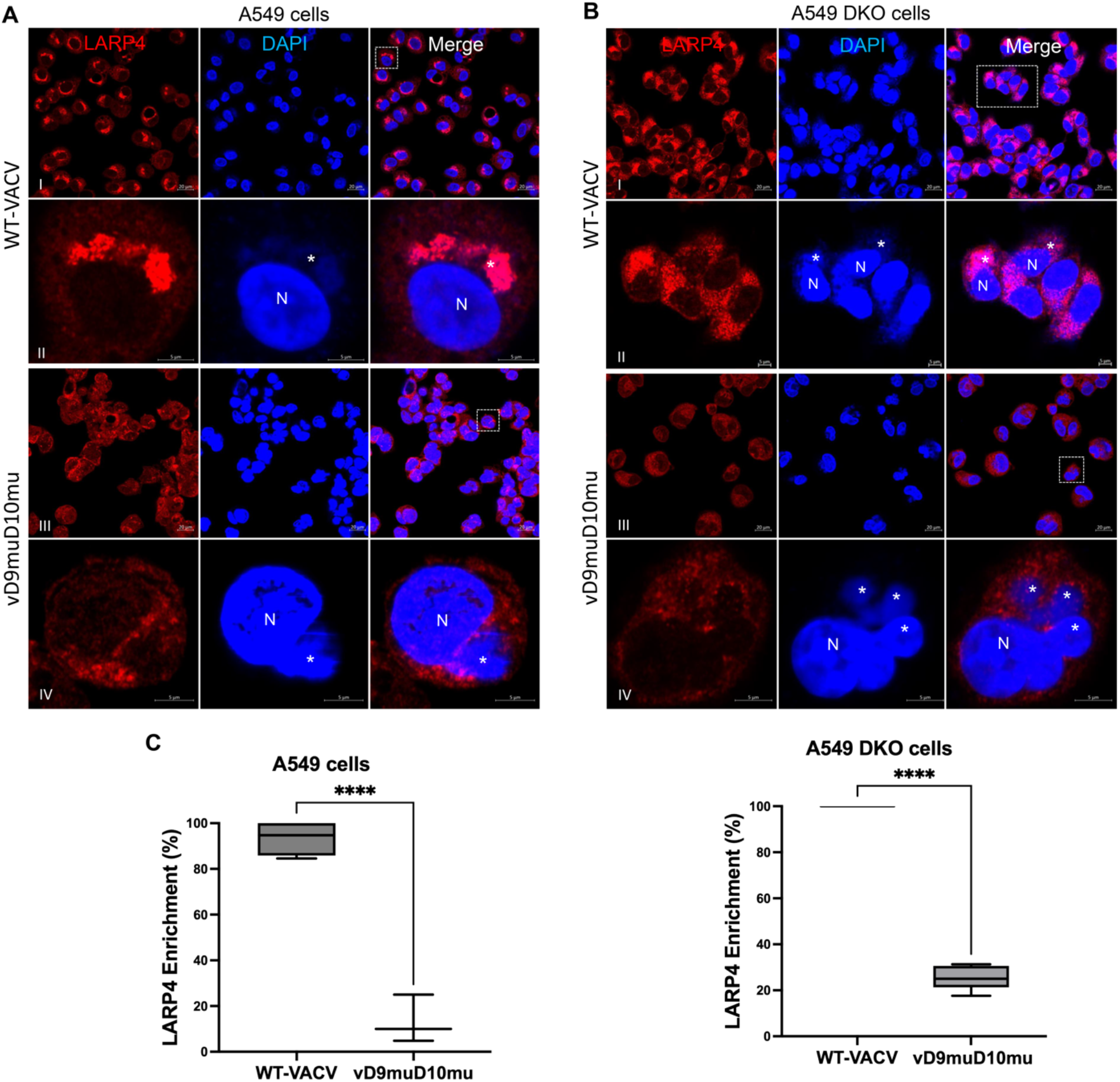
LARP4 Enrichment in viral factories requires functional VACV decapping enzymes. **(A)** Confocal immunofluorescence microscopy of WT-VACV infected A549 cells (I, II) and vD9muD10mu infected A549 cells (III, IV) (MOI=2, 12 hpi). Anti-LARP4 was used to determine the localization of LARP4 (red), and DAPI (blue) was used to stain cellular nuclei (N) and viral factories (indicated by *). (II) shows a high magnification of the cell boxed in (I), and (IV) shows a high magnification of the cells boxed in (III). **(B)** Confocal immunofluorescence microscopy of WT-VACV infected A549 DKO cells (I, II) and vD9muD10mu infected A549 DKO cells (III, IV) (MOI=2, 12 hpi). Anti-LARP4 was used to determine the localization of LARP4 (red), and DAPI (blue) was used to stain cellular nuclei (N) and viral factories (indicated by *). (II) shows a high magnification of the cell boxed in (I), and (IV) shows a high magnification of the cells boxed in (III). The presented images are the representatives of multiple fields of view. The scale bars are 5 μm and 20 μm for high and low magnifications, respectively. **(C)** Comparative analysis of LARP4 enrichment in viral factories in A549 (left) and A549 DKO (right) cells infected with WT-VACV or vD9muD10mu. Same confocal imaging specifications were used in the [aired comparison. P-values were obtained using the *Student’s t-test*, and the graph shows that LARP4 enrichment is significantly lower (P<0.01) in cells infected with vD9muD10mu than in cells infected with WT-VACV.

### Knockdown of LARP4 reduces VACV replication in HFFs

In order to investigate the role of LARP4 in VACV replication, we knocked down LARP4 expression in HFFs using two siRNAs targeting different regions of LARP4 mRNA. Successful knockdown of LARP4 protein levels was confirmed by Western blot analysis (**Fig. 5A**). The siRNA-mediated depletion of LARP4 in HFFs did not exert a noticeable decrease in nascent cellular protein synthesis using puromycin labeling (**Fig. 5A**), suggesting a negligible effect on overall cellular protein synthesis rate in HFFs. To assess VACV replication in LARP4-depleted HFFs, the cells were infected with VACV at a multiplicity of infection (MOI) of 3. Interestingly, at 12 h post-VACV infection, the HFFs with LARP4 siRNA exhibited a less severe cytopathic effect than the control siRNA-treated HFFs (**Fig. 5B**), suggesting that the LARP4 siRNA treatment may limit VACV replication. We then determined VACV titers 24 hpi by a plaque assay. Compared to the control siRNAs, LARP4 siRNAs (#1 and #2) reduced VACV titers by 10-fold and 14-fold, respectively (**Fig. 5C**). Together, these results indicate that LARP4 facilitates optimal VACV replication in HFFs.

**Figure 5.**
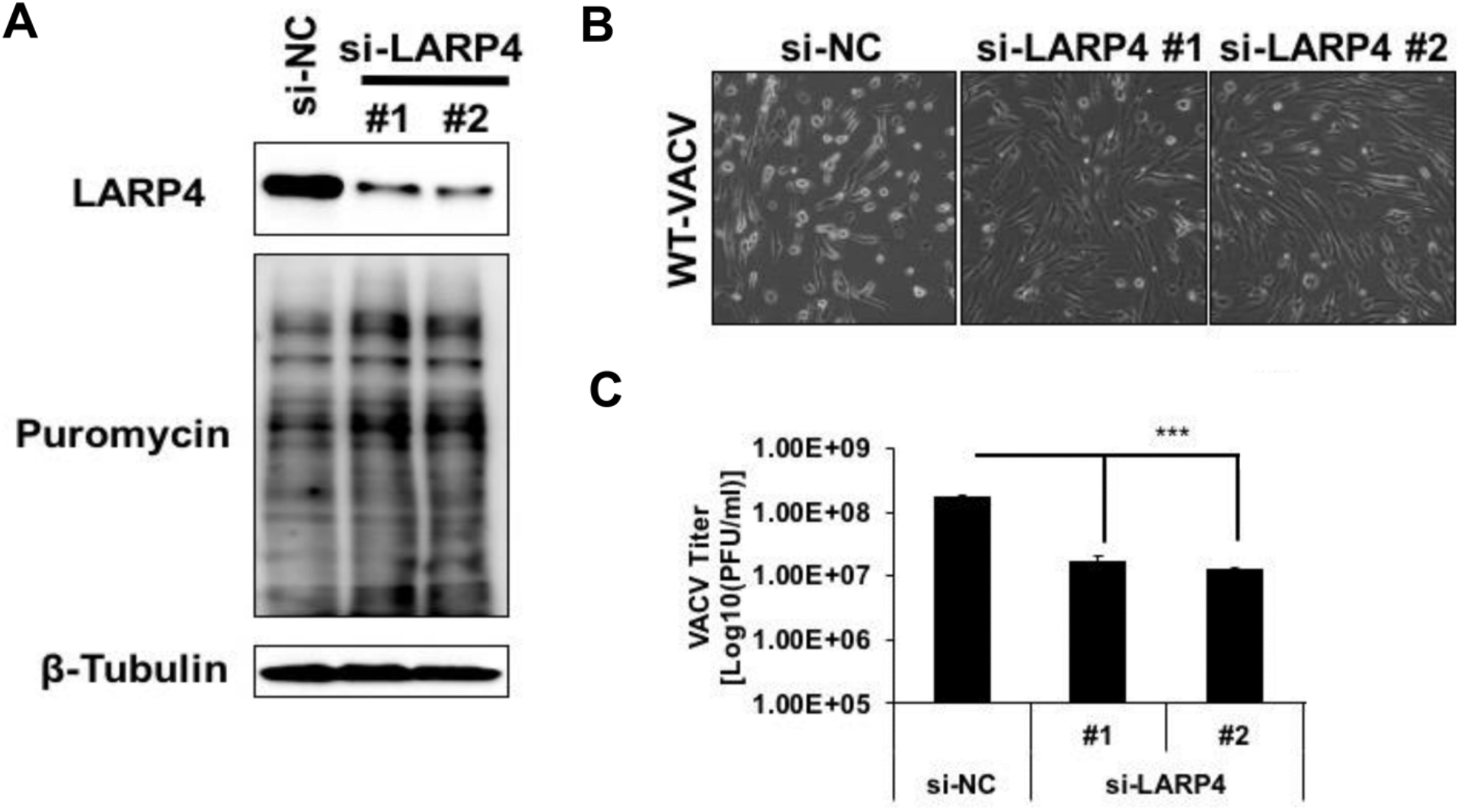
Decrease of LARP4 protein level reduces VACV replication. (**A**) HFFs were transfected with negative control siRNA (si-NC) or siRNAs targeting LARP4. Nascent protein synthesis was determined by treating cells and labeling nascent protein with puromycin (10 μg/ml) for 20 minutes at 37°C. Levels of LARP4, puromycin-labeled nascent protein, and β-Tubulin proteins were detected using specific antibodies. (**B**) At 72 hours post-siRNA transfection, HFFs were infected with VACV at an MOI of 3 VACV-infected cells were observed 12 hours post-infection by microscopy. (**C**) At 24 hpi, HFFs were collected for plaque assay. The titers of the VACV during indicated treatment were determined by plaque assay using BSC-1 cells. Error bars represent the standard deviation (SD) of at least three experiments. P-values were obtained using the *Student’s t-test;* ***P value <0.001.

### Knockdown of LARP4 has little effect on VACV early gene expression but substantially affects post-replicative gene expression

VACV replication can be divided into entry, early gene expression, viral DNA replication, intermediate and late gene expression, virion assembly, and release. We utilized Western blotting analysis to investigate the impact of LARP4 siRNA treatment on viral protein levels in HFFs. Our results demonstrated a significant decrease in viral post-replicative protein levels in LARP4 siRNA-treated cells, indicating that LARP4 is essential for viral protein synthesis. As shown in **Fig. 6A**, HFFs infected with VACV at different time points (4, 8, and 12 hpi) revealed that LARP4 siRNA treatment did not affect viral early protein E3 levels but led to a substantial reduction in the synthesis of VACV intermediate protein D13 and late protein A10. Additionally, we evaluated VACV DNA replication by quantitative PCR using DNA extracted from control and LARP4 siRNA transfected VACV-infected HFFs. AraC, an inhibitor of DNA replication, was used as a positive control. The results showed only a minor reduction (3-5-fold) in VACV DNA levels after LARP4 knockdown compared to the significant decrease after AraC treatment (453-fold decrease) (**Fig. 6B**). The findings suggest that LARP4 is primarily required for VACV post-replicative gene expression.

**Figure 6.**
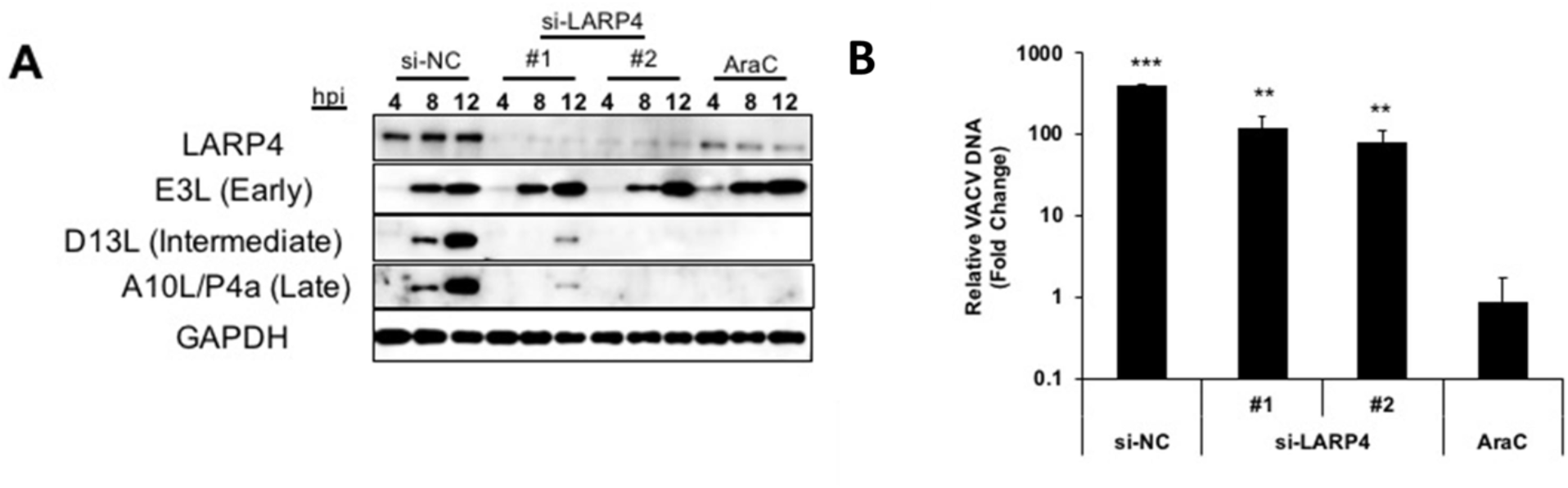
Depletion of LARP4 reduces post-replicative gene expression. **(A)** HFFs cells were transfected with negative control siRNA (si-NC) or LARP4 siRNAs. 72 hours post-transfection, HFFs were infected with VACV at an MOI of 5. AraC (40 μg/mL) treatment was used as a positive control. The HFFs were lysed, and the lysates were analyzed by Western blotting analysis to determine the levels of LARP4 at different time points (4, 8, and 12 hpi). **(B)** At 1 and 24 hpi, total DNA was extracted from the infected HFFs with indicated treatment. Levels of VACV DNA at 1 and 24 hpi were determined and represented as fold change of VACV DNA at 24 hpi to 1 hpi. Error bars represent the standard deviation (SD) of at least three experiments. P-values were obtained using *Student’s t-test;* *P value <0.05, **P value < 0.01, ***P value <0.001.

### Knockdown of LARP4 decreases 5’-poly(A) leader-mediated mRNA translation in HFFs

All VACV post-replicative mRNAs have a 5’-poly(A) leader that confers a translational advantage in VACV-infected cells [13]. LARP4 is a poly(A)-binding protein [22]. We asked if the knockdown of LARP4 affected 5’-poly(A) leader-mediated translation. Following LARP4 depletion by two siRNAs, respectively, HFFs were mock-infected or infected with VACV (MOI=5). The 5’-poly(A) leader-containing the Fluc reporter mRNA and Kozak-headed Rluc mRNA were co-transfected into uninfected and VACV-infected HFFs at 12 hpi, respectively, as described previously [13, 27]. Fluc and Rluc reporter mRNAs were either m^7^G capped (**Fig 7. AB**) or ApppG capped (**Fig 7. CD**). The translation from ApppG-capped RNA indicates cap-independent translation as this cap analog does not initiate cap-dependent translation initiation assembly [28, 29]. Relative luciferase activities of 5’-poly(A) leader-driven Fluc activities were normalized by co-transfected Kozak sequence-driven Rluc activities. LARP4 knockdown decreased translation from the m^7^G capped 5’-poly(A) leader in uninfected HFFs (**Fig. 7A**) and VACV-infected HFFs (**Fig. 7B)**. Similarly, LARP4 depletion also significantly decreased translation from ApppG-capped 5’-poly(A) leader mRNA in VACV-infected cells (**Fig. 7D**). No significant change in ApppG-capped 5’-poly(A) leader mRNA translation was observed in uninfected cells (**Fig. 7C**), which had very low Fluc expression as expected. Unlike in VACV-infected cells, AraC treatment did not alter poly(A) leader-mediated translation in uninfected HFFs (**Fig. 7A**). These results support the notion that LARP4 is required for poly(A) leader-mediated translation in both uninfected and VACV-infected cells, including cap-independent translation enhancement in VACV-infected HFFs.

**Figure 7.**
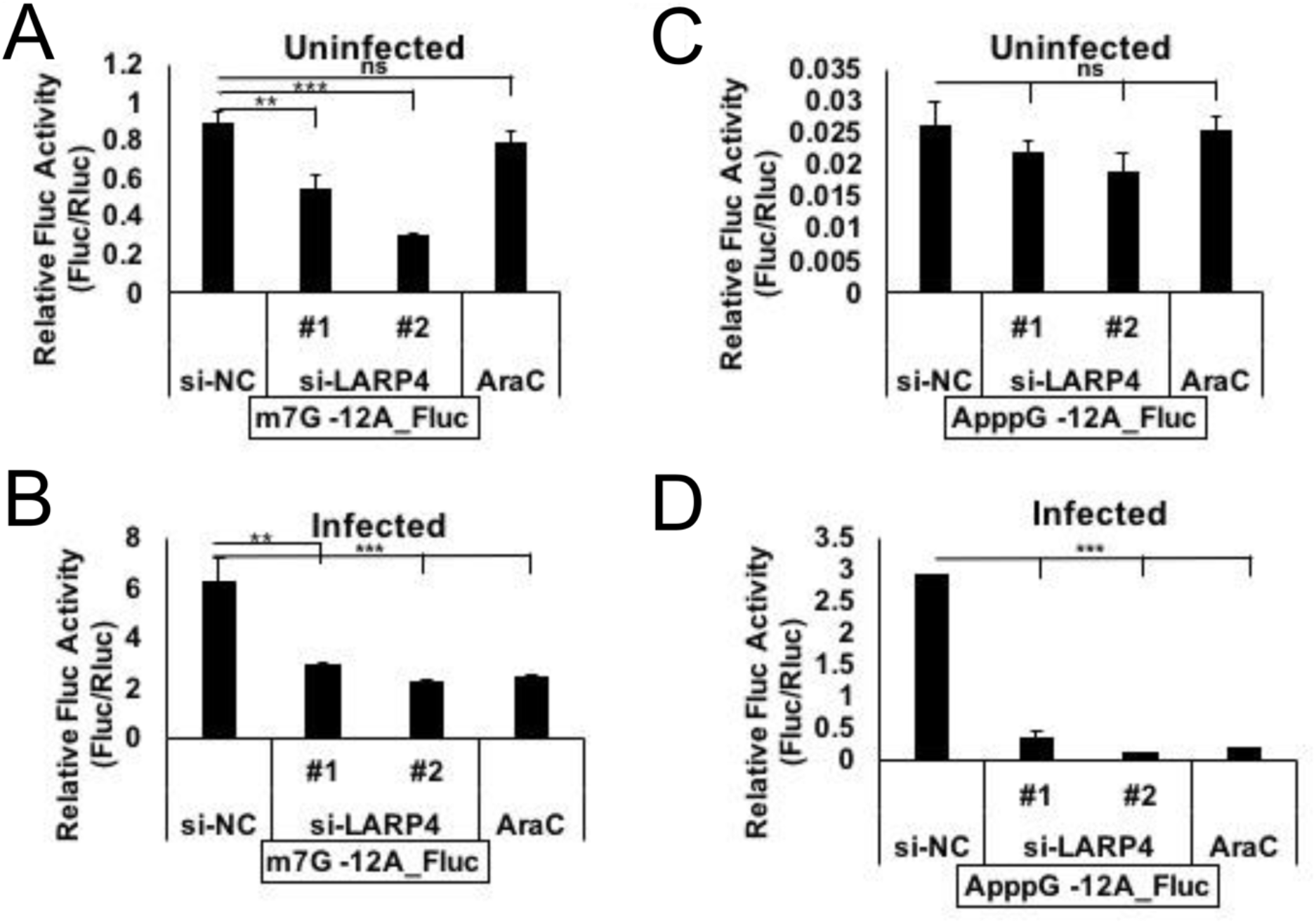
LARP4 is required for 5’poly(A) leader-mediated translation enhancement in. **HFFs.** HFFs were either infected with VACV (MOI=5) (**BD**) or kept uninfected (**AC**) after 72 hours post-transfection of control or LARP4 siRNAs. AraC (40 μg/mL) treatment was initiated after 1 hour of infection or uninfected condition. 5’-Poly(A) leader containing Fluc reporter mRNA and Kozak-headed Rluc mRNA were transfected into uninfected and VACV-infected HFFs at 12 hpi. Fluc and Rluc mRNAs were capped by either m^7^G (**AB**) or ApppG (**CD**). Relative luciferase activities were measured 5 hours after reporter mRNA transfection by normalizing 5’Poly(A) leader-driven Fluc activity by co-transfected Kozak sequence-driven Rluc activity. Error bars represent the standard deviation (SD) of at least three experiments. P-values were obtained using the *Student’s t-test;* **P value < 0.01, ***P value <0.001, ns = Not Significant (i.e., P value > 0.05).

## Discussion

Here we identified a poly(A) binding protein, LARP4, enriched in viral factories during VACV replication. Knockdown of LARP4 reduced VACV replication and 5’-poly(A) leader-mediated translation. While it has long been suspected that one or more poly(A)-binding proteins are specifically required for VACV replication, it is intriguing that the PABPC1 is excluded from the viral factories [19]. The localization of PABPC1 outside of the viral factories suggests a non-essential role of this protein in poxvirus mRNA metabolism and translation in viral factories, as viral mRNAs are produced and can be translated in the factories [19, 20]. Our data that LARP4, a poly(A)-binding protein, translocated to viral factories during VACV replication suggests the virus utilizes this specific poly(A)-binding protein to facilitate viral replication. One previous study showed that LARP4-NTD (N-terminal domain) is essential for poly(A) RNA binding (28). LARP4-NTD also contains the PAM2w motif, which is necessary for interaction with the MLLE domains of PABPC1. Cruz-Gallardo et al. demonstrated that LARP4-NTD binding is mutually exclusive with a PABP or poly(A) RNA sequence[30]. This provides a reasonable explanation for the observation that PABPC1 is excluded from the viral factories, while LARP4 is enriched in the viral factories.

Our results indicate that enrichment of LARP4 in VACV viral factories could not occur in vΔA23- and vD9muD10mu-infected cells. In vΔA23-infected cells, while viral early mRNAs, proteins are produced, and viral genomic DNA is replicated at high levels, viral intermediate mRNAs are not produced due to the deletion of one of the intermediate transcription factors A23 [24]. Subsequently, the following steps of VACV replication, including late mRNA synthesis, are also blocked. As LARP4 is a poly(A)-binding protein, this finding suggests an intermediate and late viral mRNA synthesis requirement in the factories to recruit LARP4. However, the data that LARP4 was not enriched in viral factories in vD9muD10mu-infected cells suggests the intermediate and late gene expression is not sufficient to recruit LARP4 to viral factories, as previous studies indicated that viral post-replicative gene expression, including RNA and protein synthesis occurred in vD9muD10mu-infected cells, especially in A549DKO cells [15, 25, 26]. One possible explanation for this finding is that the inactivation of the two decapping enzymes, D9 and D10, delayed and reduced the degradation of cellular mRNAs that may retain LARP4 outside the viral factories.

Our data indicate that LARP4 is required to efficiently translate mRNAs with a 5’-poly(A) leader, including cap-independent translation. While we do not have data to show how LARP4 facilitates 5’-poly(A) leader-mediated translation during VACV infection, we propose that this is due to its poly(A) binding and RACK1 binding functions [21, 22, 30]. RACK1 is a ribosomal protein of the 40S ribosomal subunit of eukaryotic cells [31], which is specifically required to translate VACV mRNAs with a 5’-poly(A) leader. During VACV infection, the S^278^ of the RACK1 STSS motif is phosphorylated, and this post-translational modification is required for poly(A) leader-mediated translation advantage [16]. One possibility is that LARP4 functions as a bridging molecule binding to the 5’-poly(A) leader and RACK1 to bring RACK1 to viral mRNAs to facilitate translation. Interestingly, RACK1 has also been found to enhance cap-independent translation from internal ribosome entry sites (IRESs) in mRNAs of several viruses, including hepatitis C virus and poliovirus [32–34]. Meanwhile, RACK1 is not essential for 5’ cap-dependent translation and cell viability [16, 32]. Our previous work has shown that the 5’-poly(A) leader is a cap-independent translation enhancer, although it is not an IRES [13]. These data support the potential role of LARP4 in promoting 5’-poly(A)-leader-mediated cap-independent translation through recruiting RACK1. RACK1 may play a broader role in cap-independent translation in addition to IRES. Additionally, VACV decapping enzymes are required to efficiently translate RNAs with a 5’-poly(A) leader [14], which is consistent with the finding in this study that LARP4 is not enriched in the viral factories in vD9muD10mu-infected cells. Additional work is required to evaluate this model.

As LARP4 is a poly(A) binding protein and promotes mRNA stability [22], it may bind VACV mRNA through the 5’-poly(A) leader and/or the 3’-poly(A) tail in the viral factories to enhance viral mRNA stability in addition to affecting RNA translation. We have previously shown that VACV decapping enzyme D10 co-localizes with mitochondria, providing a spatial mechanism for the decapping enzyme to preferentially induce the degradation of cellular mRNAs over viral mRNAs in the factories [15]. The enhancement of viral mRNA stability by LARP4 may provide another mechanism for viral mRNAs to be less likely degraded.

Knockdown of LARP4 significantly reduced VACV replication in HFFs. However, we did not observe a similar effect on VACV replication in HeLa cells with LARP4 knocked down (Data not shown). These observations suggest a cell type-specific effect of LARP4 on VACV replication, which may attribute to differential expression of LARP4 or other poly(A)-binding proteins in different cells due to their transformation statuses or tissue origins. Note that HFFs are primary cells, and HeLa cells are transformed.

Together, our data presented in this study demonstrated that LARP4 is enriched in viral factories during VACV replication. The protein is required for efficient VACV replication and translation of mRNA with a 5’-poly(A) leader in primary HFFs. The mechanism by which LARP4 influences VACV mRNA translation and other potential roles of LARP4 in viral RNA with a 5’-poly(A) leader are of great interest to pursue.

## Materials and methods

### Cell culture and virus infection

Human foreskin fibroblasts (HFFs, a gift from Dr. Nicholas Wallace) were cultured in Dulbecco’s modified eagle’s medium (DMEM, Quality Biological) containing 10% fetal bovine serum (FBS, Peak Serum) and 2 mM L-Glutamine (Quality Biological). Cells were incubated in a 5% CO2 atmosphere at 37°C. VACV Western Reserve strain (ATCC VR-1354), vD9muD10mu with both decapping enzymes inactivated, and vΔA23 with intermediate transcription factor gene A23 deleted were kindly provided by Dr. Bernard Moss and were described previously [24, 25]. Virus titer was determined by plaque assay as described elsewhere [35].

### Antibodies and chemical inhibitors

Anti-LARP4 (A5108) and anti-PABPC1 (A14872) antibodies were purchased from ABclonal Inc. Anti-GAPDH (sc-365062 HRP) antibodies were purchased from Santa Cruz Biotechnology. Anti-Puromycin (MABE343) was purchased from Sigma Aldrich. Anti-D13L and anti-A10L/P4a antibodies were gifts from Dr. Bernard Moss. The anti-E3L antibody was present from Dr. Yan Xiang. Cytosine arabinoside (AraC) was purchased from Sigma-Aldrich.

### Western blotting analysis and nascent protein analysis

Protein levels were evaluated by preparing samples as described previously [36]. The sample was resolved in the SDS-PAGE gel and transferred to a polyvinylidene difluoride (PVDF) membrane. For detection of protein, membranes were blocked in either 5% BSA or 3% Milk for 1 h at room temperature, followed by incubation with primary antibody for 1 h at room temperature or overnight at 4°C and finally, incubation with secondary antibody added in 1X TBST (with either 3% Milk or 5% BSA).

Nascent protein synthesis was determined by treating the cells with puromycin (10 μg/ml, P8833, Sigma Aldrich) for 20 min at 37°C [15]. Treatment was aborted by removing the cell culture media containing puromycin and washing once with 1xPBS. NP-40 lysis buffer was added directly to the cells, which were subsequently scrapped. Lysis was carried out by rotating the sample at 4°C for 30 min and centrifuging at 12000 X g at 4°C for 10 min. The supernatant was used for sample preparation and evaluated by western blot analysis.

### *In vitro* transcribed RNA-based luciferase assay

The RNA-based luciferase assay was used to determine the translation of 5’-Poly(A) leader containing mRNA during LARP4 depletion using the protocol described previously [13, 27]. The RNA used in this study was capped with either m^7^G (Anti-Reverse Cap Analog [ARCA], S1411L) or ApppG cap analog (S1406S, New England Biolabs).

### Knockdown using small interfering RNA (siRNA)

The small interfering RNAs (siRNAs) used for this study were purchased from Integrated DNA Technologies (IDT-DNA). Lipofectamine RNAiMax (Thermo Fisher Scientific) was used to transfect siLARP4 or siNC (Negative Control) to HFFs. The siRNAs were used at the final concentration of 12.5 nM. At 72 h post-transfection, the cells were either collected for Western blotting analysis to determine siRNA-mediated depletion of the target protein or infected with VACV for further analysis.

### Confocal Immunofluorescence assay

After desired treatment, cells were fixed with 4% formaldehyde (28908, Thermo Fisher Scientific) in 1XPBS for 20 minutes. The cells were then permeabilized with 0.1% Triton X-100 (9002-93-1, Thermo Fisher Scientific) in 1xPBS for 15 min. Cells were washed 3X with 1X-PBS and blocked for 1 h at room temperature with 2% BSA containing 1xPBS. Then, cells were incubated with the desired primary antibody (1:100 dilution in 2% BSA containing 1xPBS) for 1 h at room temperature. Again, after 3X washes, cells were incubated with a secondary antibody conjugated with a fluorophore (1:500 dilution in 2% BSA containing 1xPBS) for 1 h at room temperature. Following 2X washes with 0.1% Triton X-100 in 1X-PBS, DAPI stain (5μM) in 1xPBS was added to the cells for 20 min at room temperature. Finally, 2x washes with 1xPBS were done to remove any unbound DAPI stain. The image was taken using Carl Zeiss LSM 880 confocal microscope. ZEISS Efficient Navigation (ZEN) software was used to analyze the images.

### Quantitative Real-Time PCR

Total DNA was extracted from mock- or VACV-infected cells using E.Z.N.A. Blood DNA Kit. Relative viral DNA levels were quantified by CFX96 real-time PCR instrument (Bio-Rad) with an All-in-one 2×qPCR mix (GeneCopoeia) and primers specific for VACV genome, respectively.

## Acknowledgments

We thank Dr. Nicholas Wallace, Dr. Yan Xiang, and Dr. Bernard Moss for providing various reagents. The work was supported by the National Institutes of Health (R01AI143709 and R21AI128406) to ZY and in part by Graduate Student Summer Stipend to PD from Johnson Cancer Research Center at Kansas State University. We thank Kansas State University College of Veterinary Medicine Confocal Core for the immunofluorescence assay.

## Notes

### Competing Interest Statement

The authors have declared no competing interest.

